# Genetic deletion of cytoglobin exacerbates cardiac hypertrophy and inhibits cardiac fibroblast activation independent of changes in blood pressure

**DOI:** 10.64898/2026.01.06.697990

**Authors:** Le Gia Cat Pham, Kurrim Gilliard, Frances Jourd’heuil, Sarahann Mistretta, John J. Schwarz, Harold A. Singer, David Jourd’heuil

**Affiliations:** Department of Molecular and Cellular Physiology, Albany Medical College, Albany, NY, USA

**Keywords:** heart failure, fibroblasts, fibrosis, myofibroblasts, hemoglobins

## Abstract

Hypertension-mediated left ventricular hypertrophy and cardiac fibrosis often precede heart failure. Recent studies indicate that cytoglobin (Cygb), a globin expressed in the vasculature, increases systemic blood pressure. The present work aims to determine the role of Cygb in angiotensin II (Ang II)-induced cardiac hypertrophy and fibrosis in the mouse.

**Methods:** Males and females global Cygb knockout *(Cygb*^−/−^), and wildtype (*Cygb*^+/+^) mice were treated with Ang II (1.5 µg/kg/day) for two weeks via subcutaneous osmotic minipumps. Cardiac function was assessed through echocardiography, and hearts were analyzed for changes in hypertrophy, fibrosis, and gene expression. Functional studies were also performed in isolated cardiac fibroblasts.

**Results:** *Cygb*^−/−^ mice from both sexes showed an increase in cardiac hypertrophy over *Cygb*^+/+^ mice. Cardiac functions were also depressed in *Cygb*^−/−^ males with no changes in females. Importantly, genetic deletion of Cygb did not affect systemic blood pressure in mice, at baseline or after Ang II treatment. We established that Cygb was expressed in fibroblasts and pericytes in humans and mice hearts. Finally, we found that *Cygb*^−/−^ cardiac fibroblast did not upregulate the expression of genes associated with myofibroblasts following treatment with Ang II. This was reversed following expression of human cytoglobin.

**Conclusions:** Our findings indicate that Cygb plays a protective role in the mouse heart during Ang II-induced cardiac stress. This is the first study detailing the function of Cygb in the heart as a regulator of cardiac hypertrophy. This study also reveals a role for Cygb in regulating cardiac fibroblast activation by Ang II.

**NEW & NOTEWORTHY:** We identified cytoglobin as an important globin in cardiac pathophysiology. Genetic deletion of cytoglobin led to exacerbation of angiotensin II-mediated cardiac hypertrophy in the absence of any effect on systemic blood pressure. Cytoglobin is expressed in cardiac fibroblasts and pericytes and is required for cardiac fibroblast activation to myofibroblast. The present study reveals for the first time a role for cytoglobin in regulating angiotensin II signaling.

## Introduction

Heart failure is a leading cause of morbidity and mortality in the USA and worldwide. It is often preceded by compensatory adaptations that include ventricular remodeling characterized by cardiac hypertrophy and fibrosis. This occurs in response to acute injury such as myocardial infarction or more chronic conditions such as volume and pressure overload. In all cases, hypertension is a major risk factor for the development of heart failure and blood pressure lowering therapies are important strategies to manage or prevent heart failure.

Past studies in humans and pre-clinical models have demonstrated that hypertension is associated with increased oxidative stress and decrease in the bioavailability of the second messenger nitric oxide (NO). The latter plays an essential role in maintaining vascular homeostasis and regulating blood pressure through its vasodilatory activity and opposing hypertensive stimuli^1^. Accordingly, pharmacological inhibition of NO synthesis produces a hypertensive response^2^. In the mouse, this is not sufficient to produce ventricular remodeling, although evidence of increased interstitial cardiac fibrosis and cardiomyocyte subcellular remodeling have been documented^3^. In contrast, cardiac hypertrophy and fibrosis were clearly present in mice following pharmacological inhibition of nitric oxide synthase (NOS) combined with a high fat diet^4^. Genetic deletion of NOS3, a primary source of NO in the vasculature, reproduces the pro-hypertensive effect of pharmacological inhibition of NOS activity^5,6^. NOS3 knockout mice also develop age-dependent increase in left ventricular mass^7^ and loose cardio-protection provided by ACE inhibitors and angiotensin II type 1 receptor antagonists in heart failure after myocardial infarction^8^.

The bioavailability of NOS3-derived NO in the vasculature is regulated by multiple mechanisms. This includes oxidative inactivation to nitrate that may result from elevated superoxide production as observed in several models of hypertension^9,10^. Inactivation of NO to nitrate also occurs through reactions with hemoglobin and myoglobin and more recent studies suggest that a third globin, cytoglobin (gene code CYGB), also inactivates NO and plays an important role in regulating vascular homeostasis. Cytoglobin is expressed in the vasculature of both humans and mice, predominantly in medial smooth muscle cells and adventitial fibroblasts^11,12,13,14^. Significantly, Liu et al showed that NO consumption was attenuated in isolated aortas obtained from cytoglobin knockout mice, consistent with earlier biochemical studies demonstrating NO inactivation to nitrate by cytoglobin^15^. In the same study, global deletion of cytoglobin led to a decrease in systemic blood pressure and inhibited the pro-hypertensive effect of angiotensin II infusion. In the latter case however, the effect of cytoglobin deletion on angiotensin II-mediated cardiac remodeling was not investigated.

Based on the importance of angiotensin II as a vasopressor and the contribution of vascular cytoglobin in regulating NO bioavailability, a primary goal of the present study was to determine the extent to which cardiac remodeling following angiotensin II infusion could be decreased in mice with global deletion of cytoglobin. Surprisingly, we found that genetic deletion of cytoglobin increased cardiac hypertrophy following angiotensin II infusion in the mouse. This was accompanied with left ventricular dysfunctions in males. We found no evidence for a decrease in systemic blood pressure at baseline or following angiotensin II treatment in the cytoglobin knockout mice. In the heart, cytoglobin was primarily expressed in fibroblasts and pericytes. Our study reveals that the loss of cytoglobin altered transcriptional gene programs in hypertrophied left ventricles that were associated with decreased capillary density and inhibition of angiotensin II mediated activation of cardiac fibroblasts to myofibroblasts. *In vitro*, we show that cytoglobin is required for activation of cardiac fibroblasts to myofibroblasts.

## Methods

### Supplies and reagents

All supplies and reagents are listed in Supplementary Table S1.

### Mouse line and procedures

All experiments involving mice were approved by the Institutional Animal Care and Use Committee at Albany Medical College. Adult global wildtype and cytoglobin knockout mice were generated as previously described^13^. The blood pressure and cardiac function of 11–14-week-old mice were recorded prior to surgery. The mice were anesthetized using isoflurane (with 1–2% oxygen) before a subcutaneous pocket was created through a 1 cm mid-scapular incision. A 14-day osmotic pump (Alzet Model 1002) loaded with angiotensin-II at a dosage of 1.5 mg/kg/day or saline was inserted into the pocket. The incision was closed with sutures. At the end of the two-week infusion period, the ultrasound imaging and blood pressure were repeated prior to tissue collection. The body weight of each mouse and that of the isolated hearts were recorded as well as the length of the right femur. The heart tissue was either snap frozen in OCT media or collected into cell lysis buffer or Trizol.

Blood pressure measurements were obtained using the CODA Mouse tail cuff high-throughput acquisition system (Kent Scientific, Torrington, CT). Prior to pump insertion the mice underwent a training period of 7 days during which their blood pressure (BP) was measured daily. On the day of surgery and again just prior to tissue collection, 2 sessions of 10 BP measurements were performed for each mouse. The average of accepted readings from both sessions were used to determine the systolic, diastolic, and mean BP for each individual mouse.

To evaluate cardiac hypertrophy, cardiac structure and function were assessed using a Vevo 3100 high-resolution ultrasound imaging system (VisualSonics) before and after angiotensin II (Ang-II) treatment. Mice were anesthetized with isoflurane (1–2% in oxygen), heart rate was maintained within the physiological range throughout imaging. B-mode and M-mode echocardiographic recordings were obtained in the parasternal long- and short-axis views. Left ventricular wall thickness and chamber dimensions were measured using VevoLAB software (VisualSonics).

### Miography

Thoracic aortas were isolated from 11–14-week-old mice after anesthesia with isoflurane and euthanasia by exsanguination via vascular perfusion with ice cold physiological salt solution (PSS). The aortas were placed into a silicon bottom dissecting dish containing ice cold, carbogen aerated PSS. With the aid of a stereo microscope the perivascular adipose tissue and the tunica adventitia were gently removed, with care being taken to leave the tunica intima and tunica media layers intact. Each aorta was cut into 2 mm rings with a sharp surgical blade. Individual rings were mounted on the 200 µm pins of a DMT model 630MA myograph in aerated room temperature PSS. The mounted aortic rings were gradually warmed to 37°C and equilibrated for a minimum of 5 minutes prior to automated normalization as per the manufacturer’s directions. Using ProV8 LabChart software (ADInstruments) the isometric force was continuously recorded; responses were analyzed once the force reached a stable plateau for a minimum of 30 seconds. Washes were performed between each measurement with a minimum of four buffer changes over 20 minutes by replacing the chamber buffer with fresh aeriated PSS until the baseline returned to normal. Maximum contraction was determined by replacing the PSS with high potassium physiological salt solution (KPSS) containing 10 uM phenylephrine (PE) and recording the plateau response. The depolarization response was measured in KPSS alone. Concentration response experiments were performed with increasing concentrations of PE (1 × 10^−9^M to 3 × 10^−4^M). The relaxation response curves were obtained by pre-contracting the aortic rings with a concentration of PE corresponding to 60% of maximal PE contraction, followed by increasing concentrations of acetylcholine (Ach;1 × 10^−12^ M to 3 X 10^−6 M^). Data was analyzed with Prism software and values expressed as % of relaxation.

### Isolation, culture, and treatment of adult mouse cardiac fibroblasts

Mouse cardiac fibroblasts were isolated from 11- to 14-week-old C57BL/6 mice. Hearts were aseptically excised and rinsed twice with warm complete M199 medium (M199 supplemented with 10% fetal bovine serum, 2% Antibiotic/Antimycotic). Under sterile conditions hearts were minced into ∼1 mm² pieces, transferred to cell culture flasks with 10 ml of dissociation medium (116 mM NaCl, 2 mM HEPES, 0.94 mM NaH_2_PO_4_, 5.4 mM KCl, 5.5 mM Dextrose, 0.9 mM MgSO_4_, 0.1 mM CaCl_2_,1 mg/ml BSA, 1 mg/ml Trypsin, 133 U/ml Collagenase Type 1A, 0.02 mg/ml Pancreatin, 0.15 U/ml DNase1) and incubated at 37 °C, 5% CO₂ with gentle agitation for 20 minutes. The first 10 ml cell suspensions were transferred to 15 ml tubes; 10 ml of fresh dissociation medium were added to the undigested tissues and flasks were returned to the incubator for an additional 20 minutes. This digestion and collection process was repeated until all tissue was dissociated. The collected cell suspensions were centrifuged at 400 x g for 5 minutes and cell pellets resuspended in 1 ml of prewarmed M199 media. These cell suspensions were maintained at 37°C until the final fractions were collected then pooled and centrifuged at 400 x g for 5 minutes, the resulting cell pellet was resuspended in 4 ml of complete M199 media. Cells were plated into 6 well Tissue culture plates, and incubated under standard culture conditions for 2 hours, debris and non-adherent cells were washed away by rinsing twice with media. Adherent cells were maintained in 2 ml of fresh media for 20 hours. The plates were gently washed twice to remove dead cells, refreshed with complete medium, and returned to the incubator until reaching 80–90% confluence. These cultures were designated as passage 0 (P_0_) fibroblasts. Subsequent passages were obtained by trypsinization and reseeding in fresh complete M199. In vitro angiotensin II treatment was achieved by serum starving the cells in 0.4% FBS in M199 overnight followed by stimulation with 1 µM angiotensin II for 24 or 72 hrs.

Mouse cardiac fibroblasts (P_1_) from cytoglobin knock out mice were transiently transfected with either 1.5 µg or 3 µg of empty vector pcDNA or hCYGB following the standard DharmaFECT kb protocol for either 6 well plates (1.5 µg) or 60 mm dishes (3 µg). 24 hours post-transfection the 60 mm dishes were serum starved overnight in 0.4% DMEM with 500 µg ml geneticin. Cells were then treated with 1 µM angiotensin II for 24 hrs before collecting trizol lysates for qPCR.

### Quantitative polymerase chain reaction (qPCR)

Total RNA was extracted from tissue and cells using Trizol Reagent according to the manufacture’s protocol. Heart homogenates were prepared with a Bead Mill Homogenizer using 1.5 ml screw top tubes containing a mixed lysing matrix of 2.8- and 1.4-mm ceramic beds in 1 ml of Trizol, the mouse cardiac fibroblasts were directly lysed with Trizol. Briefly, the aqueous phase containing RNA was recovered after chloroform treatment, RNA was precipitated with isopropanol, washed with 75% ethanol, and resuspended in RNase-free water. RNA concentration and purity were assessed using a NanoDrop 2000 spectrophotometer The cDNA was synthesized using Qiagen’s QuantiTect Reverse Transcription Kits. qPCR analysis was conducted using gene-specific primers, and SsoAdvanced Universal SYBR Green super mix on a BioRad CFX Connect Realtime System equipped with CFX Maestro software. Oligonucleotide primers were designed using PrimerBLAST (NCBI) and purchased from IDT (Coralville, IA).

### Immunoblotting

All protein lysates were made in ice cold Radioimmunoprecipitation Assay (RIPA) buffer supplemented with HALT Protease and Phosphatase Inhibitor Cocktail and stored at -80°C until use. Heart homogenates were prepared with a Bead Mill Homogenizer using 1.5 ml screw top tubes containing a mixed lysing matrix of 2.8- and 1.4-mm ceramic beds with 1 ml of lysis buffer. Cultured cells were washed once with PBS and scraped directly into 150 µL lysis buffer. Once thawed the samples were spun at 20 000 x g for 20 minutes to pellet the insoluble fraction. Protein concentrations were determined using the bicinchoninic acid assay (BCA). Samples were heat-denatured in 2X or 4X Laemmli reducing sample buffer. Equal amounts of protein (10-20 µg) were loaded into corresponding wells and separated with 4–20% gradient Sodium Dodecyl Sulfate-Polyacrylamide Gel Electrophoresis (SDS-PAGE), transferred to 0.2 μm Polyvinylidene fluoride (PVDF) membranes and blocked in Tris-buffered saline containing 0.1% Tween-20 and 5% nonfat milk (TBST) overnight at 4°C. Membranes were incubated with primary antibodies for 1 h at room temperature, washed three times in TBST, and probed with the appropriate HRP-conjugated secondary antibodies for an additional 1 hour at room temperature. After 3 washes in TBST the membranes were developed with Clarity Western enhanced chemiluminescence substrate and visualized on a BioRad ChemiDoc MP imaging system with Image Lab software. Each membrane was probed a second time with either ACTB or GAPDH to serve as loading controls. The primary and secondary antibodies are listed in Supplementary Table 1.

### Masson’s trichrome staining

Masson’s Trichrome staining was used to evaluate collagen deposition and myocardial fibrosis. Fresh-frozen heart sections (10 µm) were fixed in 10% neutral buffered formalin at room temperature for 1 hour and stained using a Masson’s Trichrome Stain Kit (Polysciences Inc.) according to the manufacturer’s protocol. Briefly, sections were mordanted in Bouin’s fixative and sequentially stained with Weigert’s Iron Hematoxylin for nuclear visualization, Biebrich Scarlet-Acid Fuchsin for muscle and cytoplasm, Phosphomolybdic/Phosphotungstic acid as a differentiator, and Aniline Blue to label collagen fibers. Following staining, sections were dehydrated through 100% ethanol, cleared in xylene, and mounted with a permanent mounting medium. Images were acquired using a multimodal imaging system (BioTek Cytation 5, Agilent Technologies, Inc.) in color brightfield. Images were processed in Image J and analyzed using Color Deconvolution2 plugin.

### Hematoxylin and Eosin (H&E) staining

For general histological assessment, 10 µm frozen sections were stained with Hematoxylin and Eosin (H&E). Slides were removed from the freezer and immediately immersed in fixative solution (70% Ethanol, 4% formaldehyde, 5% glacial acidic acid) for 3 minutes. The slides were passed through 70% and 90% ethanol for 1 minute each, rinsed with tap water, and deionized water for 30 seconds each. Tissues were progressively stained in Hematoxylin for nuclear staining, rinsed, clarified and passed through bluing reagent before counterstaining with EosinY to highlight cytoplasmic and extracellular structures. The tissues were then dehydrated, cleared, and cover slipped for light microscopy.

### Immunofluorescence staining

For tissues, 10 µm thick sections were cut from prepared Optimal Cutting Temperature Imbedding Medium (OCT) blocks using a Leica CM1850 cryostat and transferred to charged microscope slides, dried at room and stored at -80°c until use. Slides were removed from the freezer, air dried, fixed with ice cold acetone for 10 minutes then dried again. After outlining the sections with a hydrophobic barrier pen tissue, sections were briefly rehydrated with PBS and blocked with 5% sera representing the secondary antibody species for at least 1 hour. The tissues were then incubated with the primary antibody or a matching isotype control diluted in blocking buffer for 1 hour at room temperature then washed three times in Phosphate Buffered Saline with 0.1% Triton X 100 (PBST). Secondary antibody was applied and incubated at room temperature for an additional hour, this step was omitted for directly conjugated primary antibodies Slides were then washed once with PBST once with a 50:50 PBS/water mix, nuclei were stained with 1 µM DAPI for 15 minutes at room temperature. After a final rinse in the 50:50 PBS/water mixture, they were cover-slipped with VectaShield Vibrance Antifade Mounting Medium, sealed with clear nail polish, and stored protected from light at 4°C.

For cellular immunofluorescence studies, cells were plated on poly-L-lysine coated 8 well glass bottom Ibidi chamber slides. Following treatment as described above, cells were fixed in 4% formaldehyde for 15 minutes at room temperature washed with PBS then permeabilized in PBS/0.2% Triton-X100 for 5 minutes at room temperature, blocked with 5% serum representing the secondary antibody for at least 1 hr. Primary antibodies and matching isotype controls diluted in blocking buffer were applied for 1 hour at room temperature, slides were washed again in PBST, incubated with secondary antibodies for an additional hour at room temperature, this step was omitted for directly conjugated primary antibodies. Slides were then washed once with PBST, once with a 50:50 PBS/water mix, nuclei were stained with 1 µM DAPI for 15 minutes. After one final wash in the 50:50 mixture, the slides were cover-slipped using VectaShield Vibrance Antifade Mounting media, sealed with clear nail polish and stored at 4°C in the dark until imaging.

Fluorescence in situ hybridization staining was performed using Advanced Cell Diagnostics RNAscope^TM^ Multiplex Fluorescent V2 Assay following the manufacturer’s protocol. For dermatopontin and cytoglobin, we used the Mm-Dpt and Mm-Cytoglobin probes, respectively. Mm-Ppib served as positive control and DapB as our negative control probe. Upon completion of this in situ hybridization protocol, sections were then stained for cytoglobin (as above).

### Bulk RNA sequencing

Total RNA was isolated from left ventricular tissue as outlined in the qPCR method above. The RNA samples were sent to GENEWIZ (Azenta Life Sciences) for sequencing. RNA integrity was evaluated by Agilent TapeStation only samples with RNA integrity number (RIN) ≥ 7.0 were used for sequencing. Differential gene expression analysis was conducted using DESeq2 in R. Genes with a P-value < 0.01 and absolute log_2_ fold change > 0.26 were considered significantly differentially expressed. Gene Ontology (GO) enrichment analysis was performed using Metascape. Custom gene set enrichment analysis was performed using a curated gene set related to myofibroblast differentiation. The gene list was assembled by integrating Gene Ontology annotations from the AmiGO2 database (Gene Ontology term myofibroblast differentiation; GO:0036446) with additional genes identified from prior literature and publicly available datasets. The final custom gene set was used for enrichment analysis against ranked gene expression data, and enrichment significance was assessed using standard permutation-based approaches.

### Analysis of single-cell RNA-Seq data

Publicly available single-cell RNA sequencing (scRNA-seq) datasets from adult mouse and human hearts were reanalyzed to determine the expression profile of cytoglobin across major cardiac cell populations. Mouse heart data were obtained from Li et al.^21^ (raw sequence data available from the Genome Sequence Archive in BIG Data Center (http://bigd.big.ac.cn/) with the accession code CRA007245), McLellan et al.^16^ (Raw sequence reads were accessed in ArrayExpress with accession number E-MTAB-8810), and human heart data were obtained from Koenig et al.^20^ (data are available on the Gene Expression Omnibus (GSE183852). Raw or processed expression matrices and corresponding metadata (cell barcodes, cluster annotations, and sample identifiers) were imported into Seurat (v4.3.0) in R (v4.2.0) for downstream analysis. Quality control filtering, normalization (LogNormalize), scaling, and principal component analysis (PCA) were performed following standard Seurat pipelines. UMAP (Uniform Manifold Approximation and Projection) was used for dimensionality reduction and visualization of cell clusters. Cell-type identities were assigned based on original annotations provided by the data source or by canonical marker gene expression. Dot plots were generated to visualize the expression of Cygb (mouse) or CYGB (human) across annotated cardiac cell types. Dot size indicates the proportion of cells expressing the gene, and color intensity reflects the average expression level in each cluster. All plots and statistics were generated in R using Seurat and ggplot2 packages.

### Statistical analysis

Statistical analyses were performed with GraphPad Prism 10.6.1. The statistical test used to analyze each data set is specified in individual figure legends and p-values are shown in figures. A p-value of less than 0.05 was considered statistically significant. Results are expressed as mean ± SEM, and statistical analysis using unpaired t-tests, one- or two-way ANOVA for two or more groups comparison followed by Tukey’s multiple comparison test were used. The number of independent replicates is also indicated in the figure legends, when applicable.

## Results

### Genetic deletion of cytoglobin increases cardiac hypertrophy in the mouse angiotensin II infusion model

To establish the effect of cytoglobin on angiotensin II-mediated cardiac hypertrophy, angiotensin II (1.5 mg/kg/day) was infused subcutaneously in 11-14 weeks female and male cytoglobin wild-type (*Cygb*^+/+^) or knockout (*Cygb*^−/−^) mice for 14 days using osmotic minipumps. We verified the absence of cytoglobin in *Cygb*^−/−^ mice in hearts by Western blotting (**Figure 1A**) and baseline echocardiography indicated no differences in heart rate, ejection fraction (EF), or fractional shortening (FS) between genotypes in either sex (**Supplementary Figure S1**). Next, heart sections were stained with Wheat Germ Agglutinin (WGA) to visualize cardiomyocyte boundaries and quantify cellular hypertrophy. We found that cytoglobin deletion significantly increased cardiomyocyte size in both sexes over the increase observed in the wild-type mice following infusion of angiotensin II (**Figure 1B and C**). Heart weight-to-body weight (HW/BW) ratios were significantly elevated in male *Cygb*^−/−^ mice after angiotensin II infusion compared to *Cygb*^+/+^ males (**Figure 1D)**. In contrast, there were no significant differences in females (**Figure 1D)**. The increased cardiac hypertrophy observed in *Cygb*^−/−^males coincided with a decrease in cardiac functions including a significant reduction in ejection fraction and fractional shortening (**Figure 1E and F**). Lastly, we found no difference in fibrosis area between *Cygb*^+/+^ and *Cygb*^−/−^mice based on Masson’s Trichrome staining (**Figure 2A and B**). Overall, these results indicate that global deletion of cytoglobin promotes cardiac hypertrophy in response to angiotensin II infusion in mice in both sexes and cardiac dysfunction in males, in the absence of additional increase in cardiac fibrosis.

**Figure 1.**
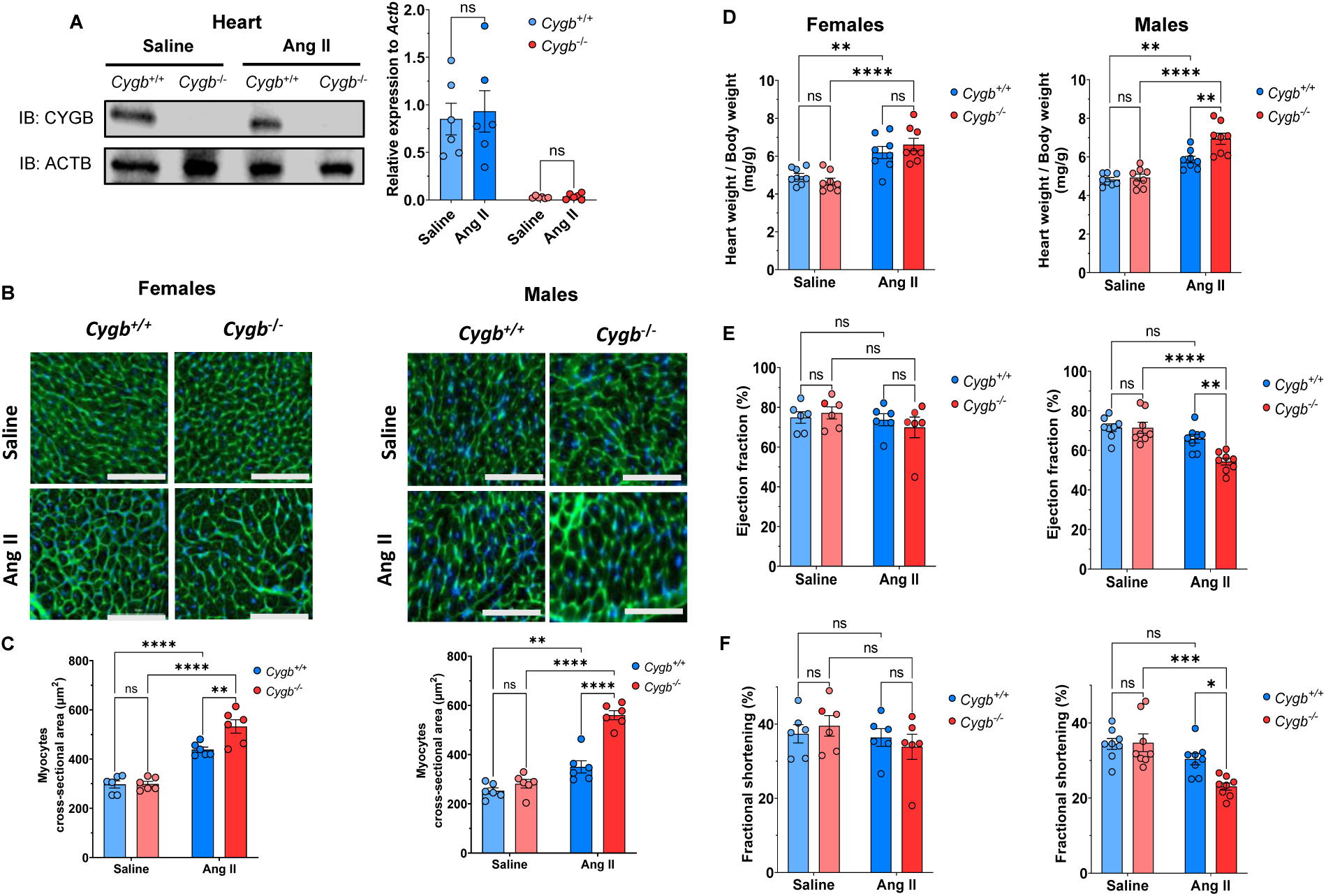
Increased cardiac hypertrophy in global cytoglobin knockout mice following angiotensin II infusion. Male and female *Cygb*^+/+^ and *Cygb*^−/−^ mice (11–14 weeks old) were infused with angiotensin II (1.5 mg/kg/day) for 2 weeks using subcutaneously implanted osmotic minipumps. After treatment, mice were euthanized, and hearts were collected for cytoglobin protein analysis by Western blotting. **A:** Left panel, representative immunoblot showing cytoglobin (CYGB) protein expression in hearts from each genotype and treatment group. ACTB (β-actin) was used as a loading control. Right panel, quantification of CYGB protein expression normalized to ACTB. **B:** Representative images of heart sections obtained from female (left panel) and male (right panel) mice and stained with wheat germ agglutinin (WGA). Scale bar: 100 µm. **C:** Quantification of cardiomyocyte cross-sectional area in the left ventricular posterior wall of hearts obtained from female (left panel) and male (right panel) hearts. **D:** Heart weight-to-body weight ratio for female (left pane) and male (right panel) mice. **E and F:** ejection fraction (E) and fractional shortening (F) were measured by echocardiography prior to sacrifice. Across the figure, each data point represents one mouse. Data are presented as mean ± SEM. Statistical analysis was performed using two-way ANOVA. *P < 0.05; **P < 0.01; ***P < 0.001, ****P < 0.0001, and ns = not significant.

**Figure 2.**
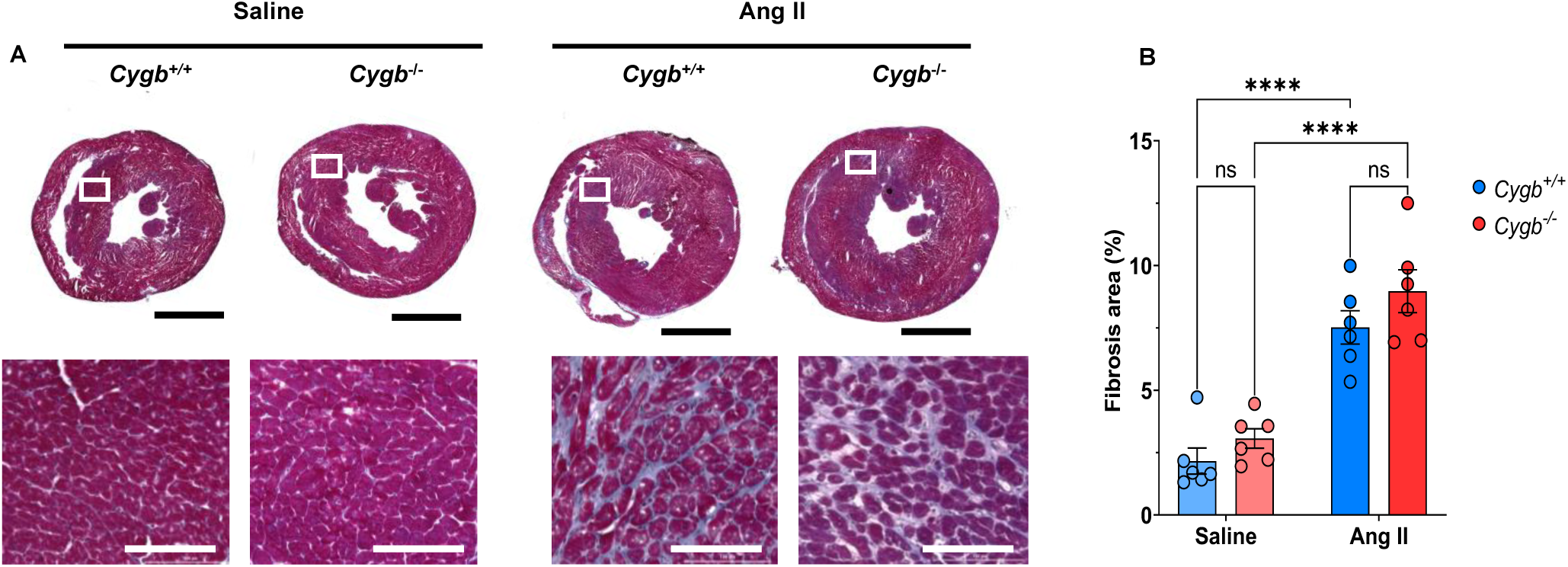
Cytoglobin deletion does not change cardiac fibrosis. **A:** Top panels, representative trichrome-stained cardiac sections taken from the shorter axes of the hearts from both *Cygb*^+/+^ and *Cygb*^−/−^ mice following infusion with either saline or angiotensin II (Ang II), as described in Figure 1. Scale bars are 2000 µm. Bottom panels are enlargements of white insets **B:** Changes in cardiac fibrosis expressed as a percentage of total area in male mice with and without Ang II infusion. Scale bar = 100 µm. Across the figure, each data point represents one mouse. Data are presented as mean ± SEM, with statistical significance determined using two-way ANOVA. *P < 0.05, **P < 0.01; ***P < 0.001, ****P < 0.0001 ns = not significant.

### Systemic blood pressure is not altered in the cytoglobin knockout mice at baseline or following Angiotensin II infusion

To determine whether the changes in cardiac hypertrophy and function observed in cytoglobin knockout mice were secondary to alterations in systemic hemodynamics, we assessed blood pressure and vascular functions. At baseline, systolic, diastolic, and mean arterial pressures were not different between genotypes (**Figure 3A and Supplementary Figure S2**). Similarly, acetylcholine (ACh)-induced endothelium-dependent vasorelaxation in isolated aortic rings was not different between groups (**Figure 3B**). Following two weeks of Ang II infusion, both genotypes exhibited the expected increase in blood pressure. However, no significant differences in systolic, diastolic, or mean arterial pressures were observed between *Cygb*^+/+^and *Cygb*^−/−^mice (**Figure 3C and Supplementary Figure S2**). These findings indicate that the exacerbated hypertrophy and systolic dysfunction observed in *Cygb*^−/−^mice occur independently of additional changes in systemic blood pressure or vascular reactivity.

**Figure 3.**
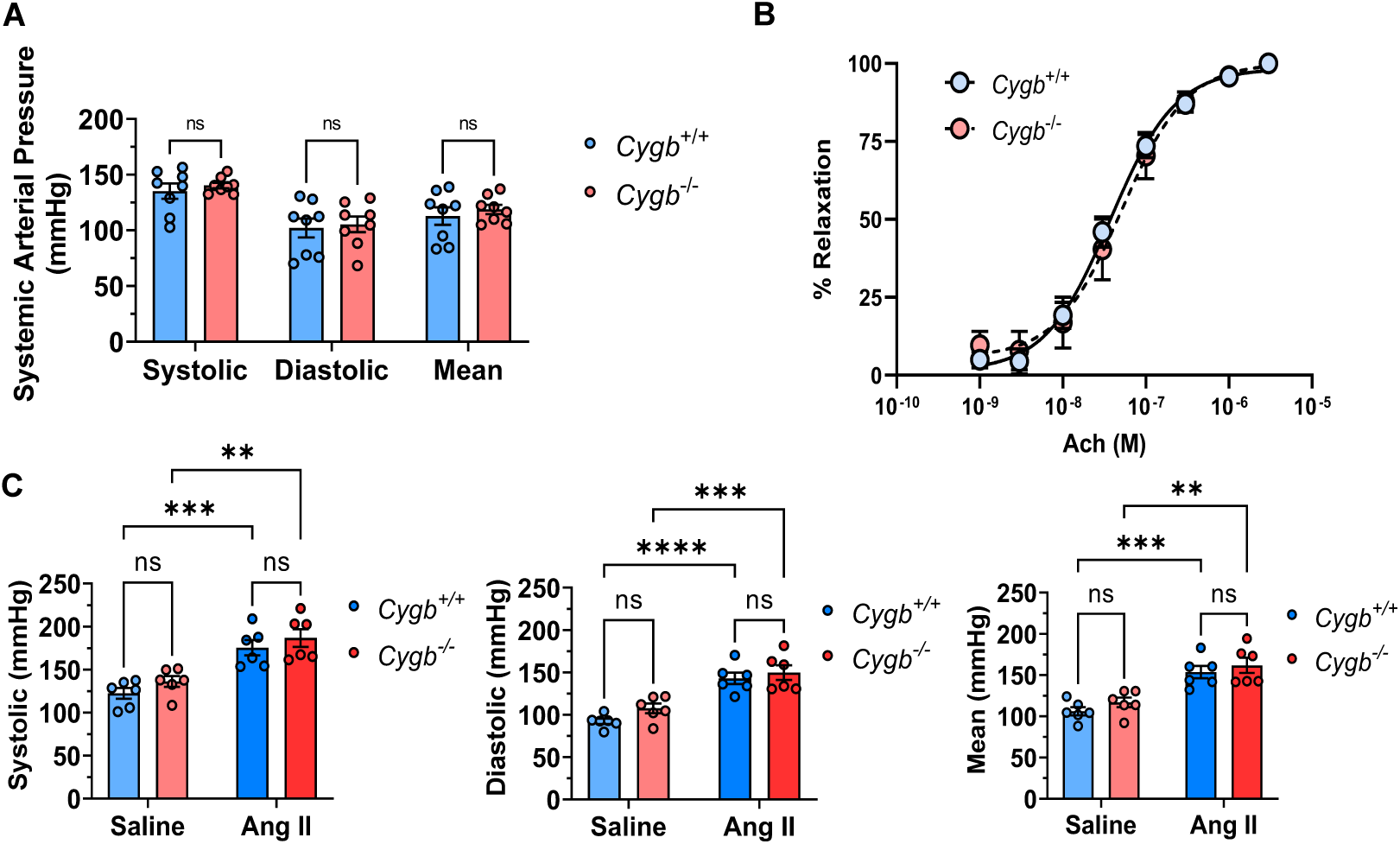
Genetic deletion of cytoglobin does not alter blood pressure at baseline or following Ang II infusion. **A:** baseline blood pressure in male *Cygb*^+/+^ and *Cygb*^−/−^ mice measured with a tail-cuff system. **B:** Acetylcholine (ACh)-induced vasorelaxation in isolated aortic rings (n = 4). **C:** Male *Cygb*^+/+^ and *Cygb*^−/−^ mice (11–14 weeks old) were infused with saline or angiotensin II (Ang II; 1.5 mg/kg/day) for 2 weeks via subcutaneously implanted osmotic pumps. Systolic, diastolic, and mean arterial blood pressures were measured with a tail-cuff system. Data are presented as mean ± SEM, with statistical significance determined using two-way ANOVA. **P < 0.01; ***P < 0.001, ****P < 0.0001, ns = not significant.

### Cytoglobin is expressed in cardiac fibroblasts and pericytes

Previous work indicated expression of cytoglobin in the heart and association with cardiac fibroblasts^17,18^ and cardiomyocytes^19^. Using a publicly available human^20^ (Gene Expression Omnibus GSE183852) and mouse^21^ (accession code CRA007245) single-cell RNA sequencing dataset, we found that cytoglobin mRNA transcripts were enriched in fibroblasts and pericytes in humans and mice and below detection in other cardiac cells including cardiomyocytes and endothelial cells (**Figure 4A and B**). This was further validated with a second mouse heart dataset that included control and angiotensin II-treated groups^16^ (accession number E-MTAB-8810; **Supplementary Figure S3)**. Cytoglobin transcripts co-associated with cell clusters expressing the fibroblast marker dermatopontin (*Dpt*) and pericyte marker neuron glial antigen 2 (*Ng2*; **Supplementary Figure S3A**). There were no evident changes in cytoglobin expression following angiotensin II infusion, consistent with quantification of cytoglobin protein content by Western blotting (**Supplementary Figure S3B and Figure 1**). Unsupervised clustering to identify fibroblast subpopulations revealed broad expression of cytoglobin in fibroblasts across all subclusters **(Supplementary Figure S3C and D)**. Interestingly, cytoglobin was also expressed in thrombospondin-4 (Thbs4) positive fibroblasts that are more abundant following angiotensin II infusion^16^ **(Supplementary Figure S3D)**.

**Figure 4.**
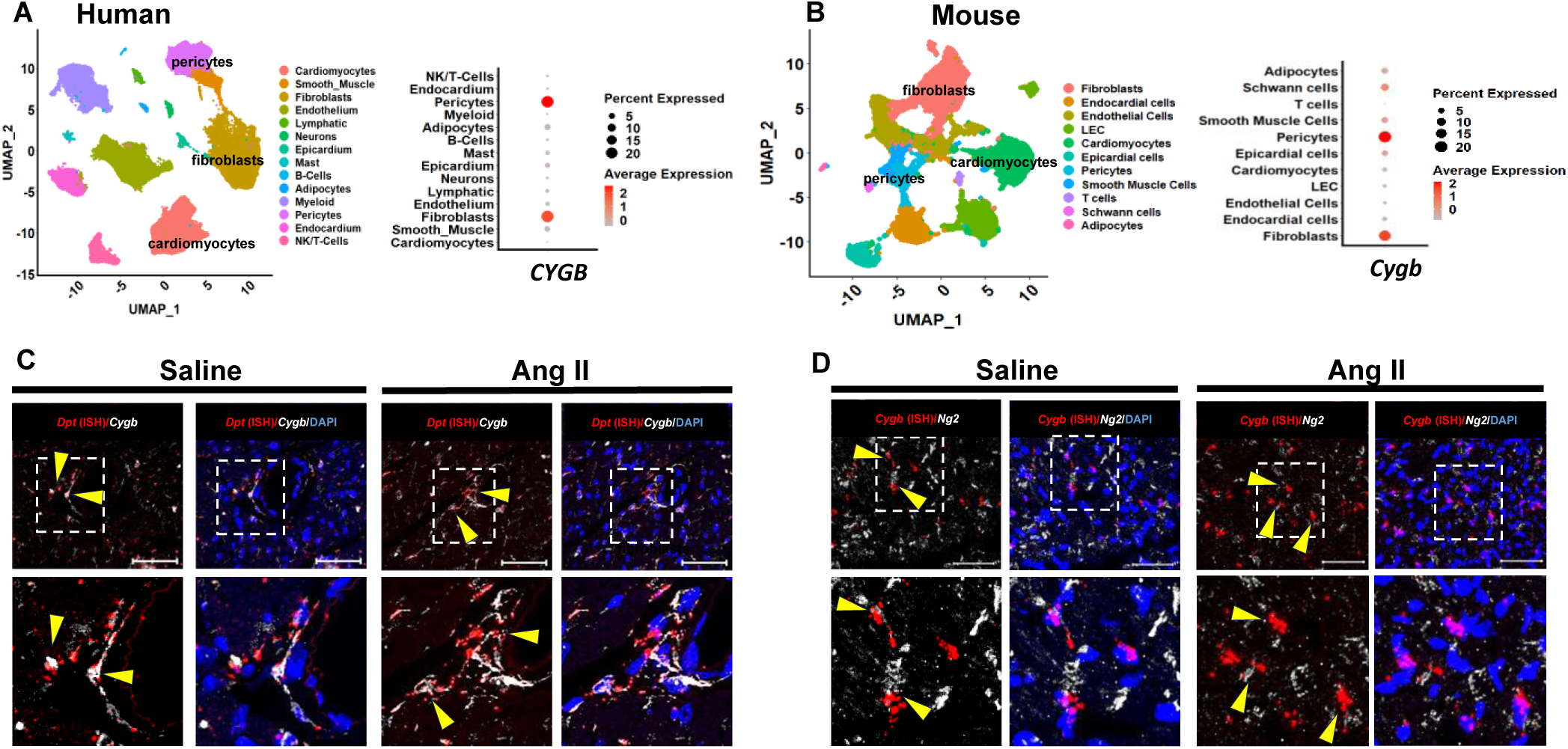
Cytoglobin is expressed in cardiac fibroblasts and pericytes. **A-B:** Analysis of publicly available single cell RNA-seq data sets for human (GSE183852) and mouse (CRA007245) hearts. For A and B, left panel is the Uniform Manifold Approximation and Projection (UMAP) of isolated cardiac cells. The right panel is the dot plot of cytoglobin expression in identified clusters. **C:** Cardiac tissue sections were prepared for *in situ* hybridization for the fibroblast marker dermatopontin (*Dpt*, red) and immunostaining for cytoglobin (CYGB, gray) with DAPI as a nuclear stain. Representative immunofluorescence staining are shown for wild-type male mice following 2-week infusion of saline or Ang II.

Next, immunostaining with an antibody against cytoglobin combined with *in situ* hybridization for dermatopontin mRNA confirmed expression of cytoglobin in perivascular and interstitial fibroblasts at baseline and following angiotensin II infusion (**Figure 4C**). Similarly, co-association was evident between NG2 (gene code *Cspg4*) protein and cytoglobin mRNA visualized by *in situ* hybridization, consistent with cytoglobin expression in pericytes (**Figure 4D**). Cardiac pericytes are vascular mural cells that are structurally and functionally associated with heart capillaries^22,23,24^. Because changes in capillary density may determine the myocardial response to chronic pressure overload^25,26,27^ and since we show that cardiac pericytes express cytoglobin, we next determined whether cytoglobin deletion altered heart capillarization. There was no difference in capillary density at baseline between the wildtype and knockout mice, based on CD31 immunostaining of left-ventricular sections (**Supplementary Figure 3**). However, there was a small but significant decrease in capillary density in the cytoglobin knockout mice over the decrease observed in the wildtype mice following angiotensin II infusion (**Supplementary Figure S4**).

### Cytoglobin is required for cardiac fibroblast activation by angiotensin II

To obtain additional mechanistic insights, we performed bulk RNA sequencing of left ventricular heart tissue from male *Cygb*^+/+^ and *Cygb*^−/−^ mice following 2 weeks of angiotensin II infusion. Differential gene expression analysis identified a distinct transcriptional signature associated with cytoglobin deletion, with 548 genes significantly upregulated and 327 genes downregulated in *Cygb*^−/−^ mice (**Figure 5A**). Gene ontology analysis of upregulated transcripts revealed enrichment for biological processes related to cardiac conduction, cellular response to interferon β, and DNA damage pathways (**Figure 5B**). Downregulated genes were enriched for genes associated with blood vessel development and extracellular matrix organization (**Figure 5B and C**). The gene list corresponding to the biological process “Extracellular matrix organization” included genes such as *Sox9*, *Acta2*, *Eln*, and *Comp*, which are associated with the activation of fibroblasts to myofibroblasts (**Figure 5C**). To further explore the role of cytoglobin in cardiac fibroblast activation, we performed a gene set enrichment analysis^28,29^ for genes associated with myofibroblast differentiation. We used a gene list compiled from different sources focusing on genes that represent important myofibroblast markers^30,31,32,33,34,35,36,37^ (**Figure 5D**). With a normalized enrichment score of -1.9684 and an FDR q value <0.0001, the analysis results indicated a strong association between cytoglobin deletion and inhibition of cardiac fibroblast differentiation to myofibroblasts (**Figure 5D**).

**Figure 5.**
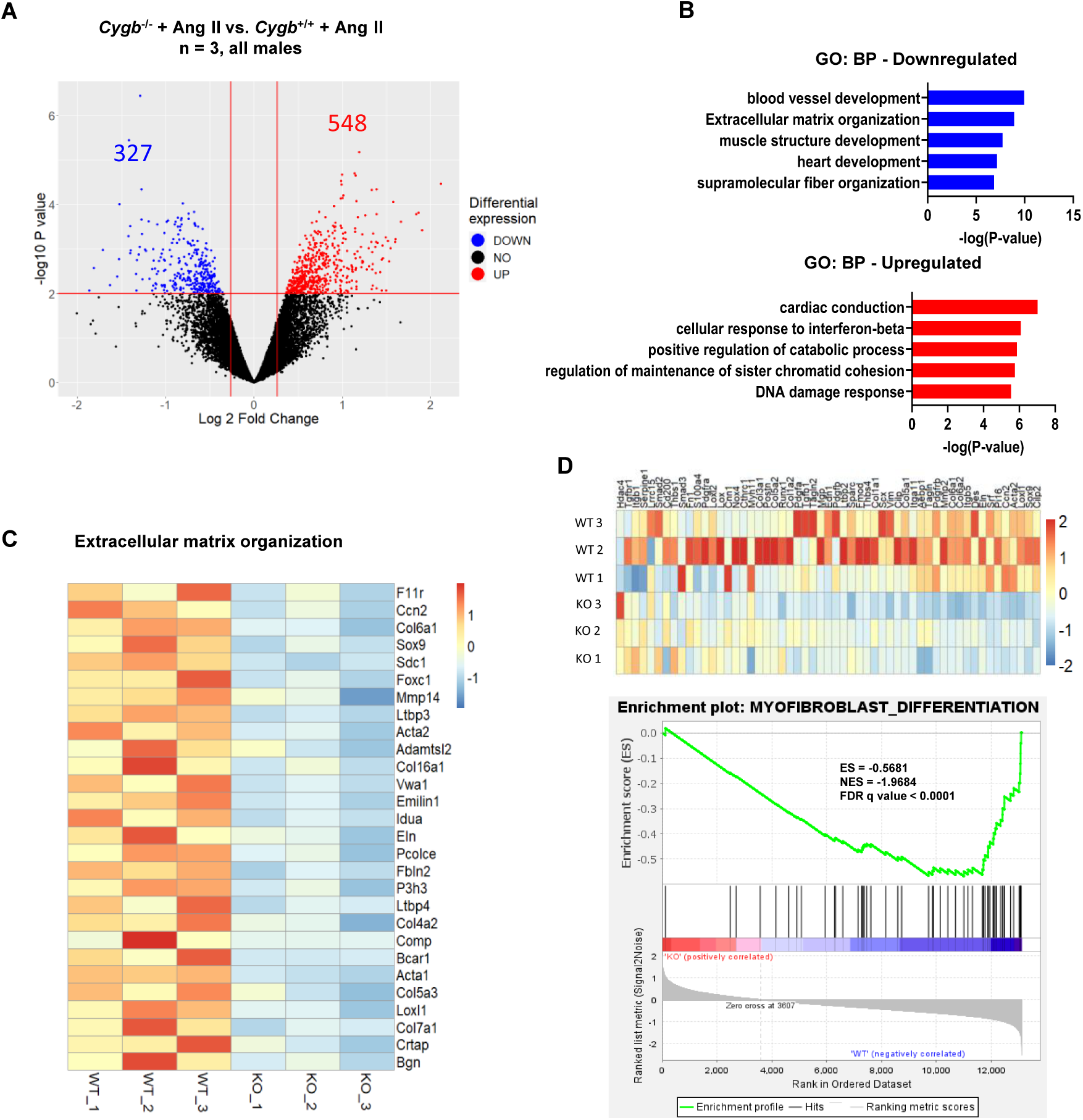
RNA seq analysis reveals a role for cytoglobin in myofibroblast dedifferentiation. Male *Cygb*^−/−^ and *Cygb*^+/+^ mice (n = 3 per group) were infused with angiotensin II (Ang II; 1.5 mg/kg/day) for 2 weeks. A: Volcano plot showing differentially expressed genes between genotypes; numbers indicate upregulated (red) and downregulated (blue) transcripts in *Cygb*^−/−^hearts. B: Top Gene Ontology biological processes (GO:BP) enriched among downregulated (top panel) and upregulated (bottom panel) genes. C: Heatmap of genes associated with extracellular matrix organization pathway. Data were analyzed using DESeq2 for differential expression, followed by GO enrichment analysis and GSEA for pathway-level interpretation. D: Heat map and GSEA enrichment plot of the gene set for myofibroblast differentiation.

To investigate whether cytoglobin has direct effects on cardiac fibroblast activation, we isolated adult cardiac fibroblasts from cytoglobin *Cygb*^+/+^ and *Cygb*^−/−^ mice. Prior to passage (P_0_ cells), indirect immunofluorescence staining for cytoglobin indicated that cytoglobin was expressed across the cell preparation and that more than 70% of the cells expressed the fibroblast marker dermatopontin (*Dpt*; **Figure 6A**). Similar to previous studies on freshly isolated medial vascular smooth muscle cells, we found that cytoglobin protein levels were decreased following passage of the cardiac fibroblasts^12^ (**Figure 6B**). Next, P_1_ cardiac fibroblasts from *Cygb*^+/+^ and *Cygb*^−/−^ mice were treated with either saline or angiotensin II. Quantitative real time polymerase chain reaction confirmed the deletion of cytoglobin in the *Cygb*^−/−^ cells (**Figure 6C**). Most notably, the angiotensin II-mediated activation of cardiac fibroblasts to myofibroblasts was inhibited in *Cygb*^−/−^ cells as revealed by the absence of *Acta2*, *Postn*, and *Col1a1* upregulation (**Figure 6C**) and the decrease in ACTA2 positive cells determined by immunofluorescence (**Figure 6D**). The angiotensin II-stimulated expression of the fibrogenic mediators CTGF (*Ccn2*) and transforming growth factor beta (TGFβ) was also inhibited in *Cygb*^−/−^ cells (**Figure 6C**). Finally, to ascertain a causative link between cytoglobin and fibroblast activation, we performed a rescue of function experiment by transfection of human cytoglobin in *Cygb*^−/−^ cardiac fibroblasts followed by angiotensin II treatment. Expression of human cytoglobin (hCYGB) in *Cygb*^−/−^ cardiac fibroblasts was confirmed by Western blot and qRT-PCR (**Figure 6E and F**) and we measured the relative expression of *Acta2, Col1a1, Postn, Ccn2, and Tgfb1*mRNA transcripts by qRT-PCR (**Figure 6F**). There was no difference in transcript levels between cells expressing empty vector and hCYGB when treated with PBS. However, all the transcripts measured were significantly increased in *Cygb*^−/−^ cells expressing hCYGB when treated with angiotensin II. These results revealed a direct causative link between cytoglobin and angiotensin II-mediated cardiac fibroblast activation.

**Figure 6.**
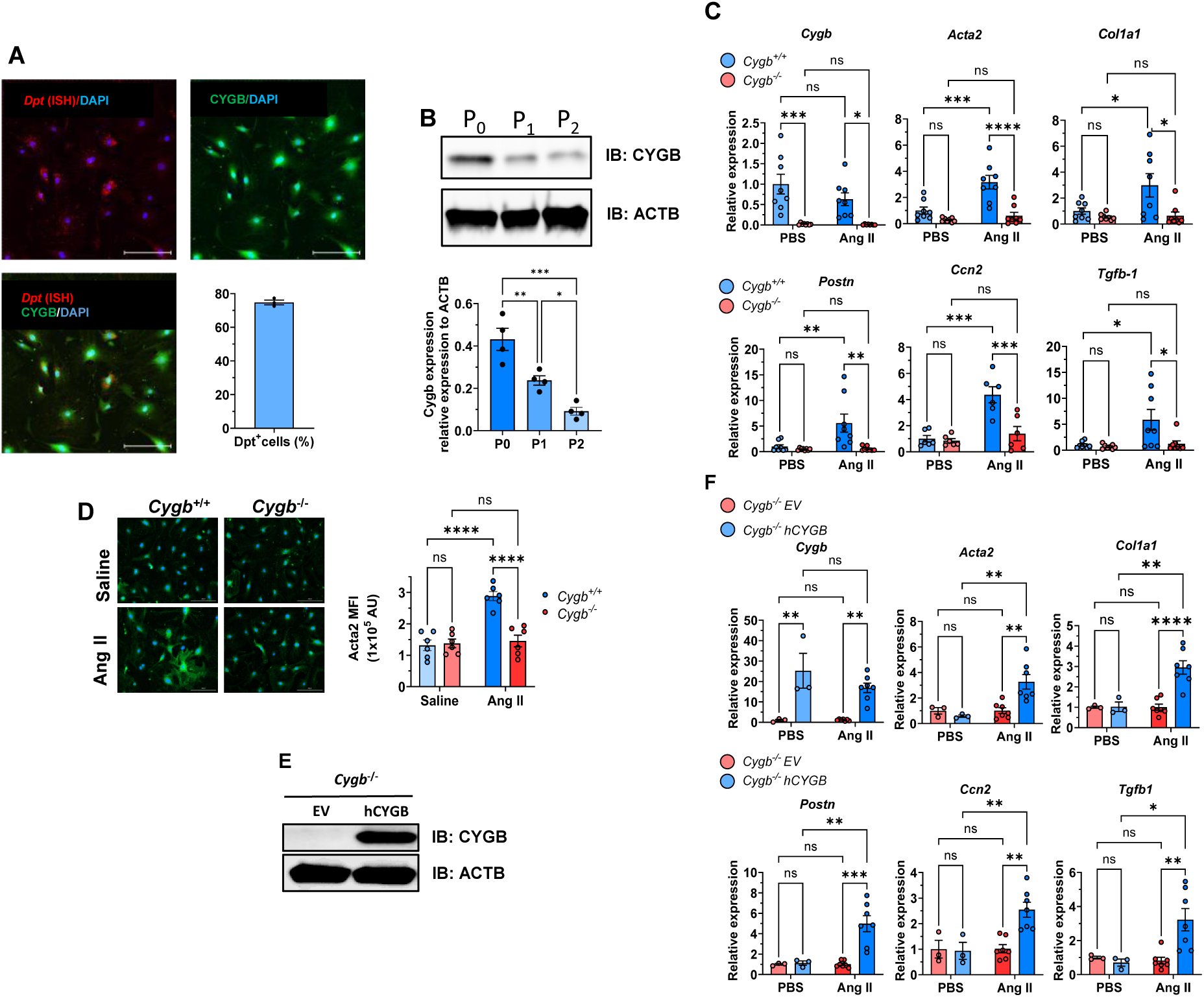
Cytoglobin is required for angiotensin II-mediated cardiac fibroblast differentiation to myofibroblast *in vitro*. **A:** Cardiac fibroblasts were isolated from wildtype mice. *In situ* hybridization for dermatopontin (Dpt, red) combined with indirect immunostaining for cytoglobin (CYGB, green) and nuclear stain (DAPI, blue) was performed and visualized by confocal microscopy. Bar graph shows the number of Dpt positive cells with all cells cytoglobin positive. **B:** Cell lysates were prepared following initial isolation (P_0_) and after one (P_1_) or two (P_2_) passages. Cytoglobin (CYGB) expression was determined by Western blotting (top panel) using beta actin (ACTB) as an internal standard and quantitation is shown in the bottom panel. **C:** Freshly isolated *Cygb*^−/−^ and *Cygb*^+/+^ cardiac fibroblasts were treated with phosphate-buffered saline (PBS) or 1µM angiotensin II (Ang II) for 24 hours. Relative expression levels of *Cygb*, *Acta2*, *Col1a1*, *Postn*, *Ccn2*, and *Tgfb-1* genes were quantified by qRT-PCR. **D**: Top panel, representative confocal images of cardiac fibroblasts from male *Cygb*^+/+^ and *Cygb*^−/−^ mice treated with angiotensin II and stained for Acta2 with quantitation of mean fluorescent intensity on the bottom panel. **E:** Freshly isolated *Cygb*^−/−^ cardiac fibroblasts were transiently transfected with an empty vector (EV) or a vector for expression of human cytoglobin (hCYGB). Cell lysates were prepared and cytoglobin expression was determined by Western blotting using beta actin (ACTAB) as an internal standard. **F:** Freshly isolated *Cygb*^−/−^ cardiac fibroblasts were transfected with and empty vector (EV) or a vector for expression of human cytoglobin (hCYGB) and then treated with phosphate-buffered saline (PBS) or 1µM angiotensin II (Ang II) for 24 hours. Relative expression levels of *Cygb*, *Acta2*, *Col1a1*, *Postn*, *Ccn2*, and *Tgfb-1* genes were quantified by qRT-PCR. Data are presented as mean ± SEM, with statistical significance determined using two-way ANOVA. *P < 0.05, ****P < 0.0001, and ns = not significant.

## Discussion

In the present work, we found that the loss of cytoglobin exacerbates cardiac hypertrophy in mice following angiotensin II infusion. This result was unanticipated given a previous study by Zweier et al. showing that global deletion of cytoglobin in the mouse was associated with sustained decrease in systemic blood pressure and inhibition of the pro-hypertensive effect of Ang II infusion^15^. In contrast, we found no changes in systemic blood pressure, angiotensin II-induced hypertension, or acetylcholine-mediated vasorelaxation. Zweier et al. used 48-week-old mice, and it is possible that the age of the mice, in addition to their different genetic background, altered the expression and cellular distribution of cytoglobin. Previous studies have also shown an increased inflammatory burden in older cytoglobin knockout mice with increased NO production through activation of NOS2, which might explain the hemodynamic changes observed in the Zweier study. We did not pursue additional work to firmly establish that NO bioavailability and NO dependent vasoreactivity were unaltered in our cytoglobin knockout mouse line. However, our results suggest that the function of cytoglobin in the cardiovascular system cannot be explained solely based on its reaction with NO. Biochemical studies which indicate strong hydrogen peroxide scavenging activity of cytoglobin *in vitro* and *in vivo* further support the possibility of alternative functions and mechanisms of action^14^.

It was notable that although the exacerbation of cardiac hypertrophy was observed in both sexes, decrease in left ventricular function was only evident in males. As shown in **Figure 1**, the effect size on cardiomyocyte area was smaller in female knockout mice following angiotensin II infusion due to a greater increase in the female wildtype mice. Importantly, the present study was not specifically designed to address sex differences, which would require larger group size and more detailed experimental considerations related for example to the angiotensin II dosage and sampling time. Additional work is also warranted to examine age-related alterations in systemic hemodynamics and cardiac remodeling following manipulation of cytoglobin levels and pro-hypertensive stressors.

We establish that in the heart, cytoglobin is expressed primarily in fibroblasts and pericytes. Absence of cytoglobin expression in cardiomyocytes was previously noted^17,18^. However, a later study found that heart cytoglobin protein expression was increased in a mouse line with cardiomyocyte specific overexpression of activated calcineurin^19^. Although the same study characterized cytoglobin expression *in vitro* in C2C12 myocytes, specific association of cytoglobin with cardiomyocytes *in vivo* was not established. Significantly, mice with cardiomyocyte specific overexpression of activated calcineurin develop extensive interstitial fibrosis, in addition to cardiac hypertrophy^38^. Thus, it is possible that the increase in cytoglobin protein expression observed by the Mammen group in this mouse line was due to the expansion of activated fibroblasts expressing cytoglobin, rather than cardiomyocytes. Overall, our results showing expression of cytoglobin within perivascular fibroblasts of coronary arteries – in addition to interstitial fibroblasts – is consistent with our previous work showing that cytoglobin is also expressed in adventitial fibroblasts of large arteries such as the aorta and carotid arteries^11^.

Our results indicate that the loss of cytoglobin exacerbated cardiomyocyte hypertrophy with no change in fibrosis. The specific mechanism by which this may occur is unclear because of the lack of effect of cytoglobin deletion on blood pressure and the absence of cytoglobin in cardiomyocytes. The lack of effect on fibrosis is even more striking considering the inhibitory effect of cytoglobin on cardiac myofibroblast differentiation and could suggest alternative sources for extracellular matrix proteins. The increased expression of extracellular matrix related proteins in the heart following chronic infusion of angiotensin II has been clearly established. However, the demonstration that cardiac fibroblasts are primary collagen producing cells in response to angiotensin II *in vivo* is lacking. Instead, bone marrow and endothelial-to-mesenchymal derived fibrocytes have been proposed as alternative sources of extracellular matrix proteins. For example, a set of studies has established that angiotensin II infusion stimulates the production of MCP-1 by endothelial cells, that is in turn required for the accumulation of myeloid-derived collagen-producing fibrocytes^39,40,41,42,43^. Angiotensin II infusion in the mouse also increases the expression of other pro-inflammatory cytokines and growth factors including TGFβ, interleukin-1, and tumor necrosis factor, all of which have been implicated in cardiomyocyte hypertrophy^44,45,46,47,48^. Thus, it is possible that cytoglobin deletion exacerbates the inflammatory response to enhance cardiac hypertrophy. This is suggested by the upregulation of genes associated with interferon beta (**Figure 5B**) and future studies will have to establish how fibroblast and pericyte cytoglobin contributes to this process.

Lastly, our study suggests for the first time that cytoglobin is essential for the activation of cardiac fibroblasts by angiotensin II. The inhibition of smooth muscle alpha actin (ACTA2) expression – a marker of myofibroblast differentiation – in the absence of cytoglobin was reminiscent of previous work, in which the loss of ACTA2 expressed in vascular smooth muscle cells was accelerated in cytoglobin-deficient mice with carotid artery^13^. However, this was in contrast with previous work showing increased expression of ACTA2 in cytoglobin-deficient mouse hepatic stellate cells *in vitro*^49^. Overall, these and other studies support the idea that cytoglobin is an important regulator of myofibroblast differentiation^17,18,49,50^. They also indicate differences in the functional engagement of cytoglobin across cell types, which will need to be mechanistically addressed in the future. Work from this laboratory and others indicate that cytoglobin rapidly reacts with specific reactive oxygen species (ROS) such as hydrogen peroxide^14,51,52,53^. The role of ROS and NADPH oxidases in angiotensin II-dependent hypertension and remodeling is well established^54^. There is also evidence to support the role for fibroblast NADPH oxidases in regulating vascular remodeling and tissue fibrosis^55,56^. Our previous work in serum-stimulated vascular smooth muscle cells suggested a link between cytoglobin, NOX4, and gene expression through interaction with chromatin remodelers such as HMGB2^13^. We would like to propose that the molecular function of fibroblast cytoglobin directly intersects with angiotensin II signaling by regulating redox signals either proximal to the angiotensin II receptor or more distal, through redox regulation of angiotensin II responsive transcription factors.

In summary, we established that cytoglobin inhibits cardiac hypertrophy independent of blood pressure following Ang II infusion in the mouse. To our knowledge, this is the first study indicating a role for cytoglobin in regulating cardiac hypertrophy and angiotensin II signaling.

## Supplementary Figures

**Supplementary Table S1.**
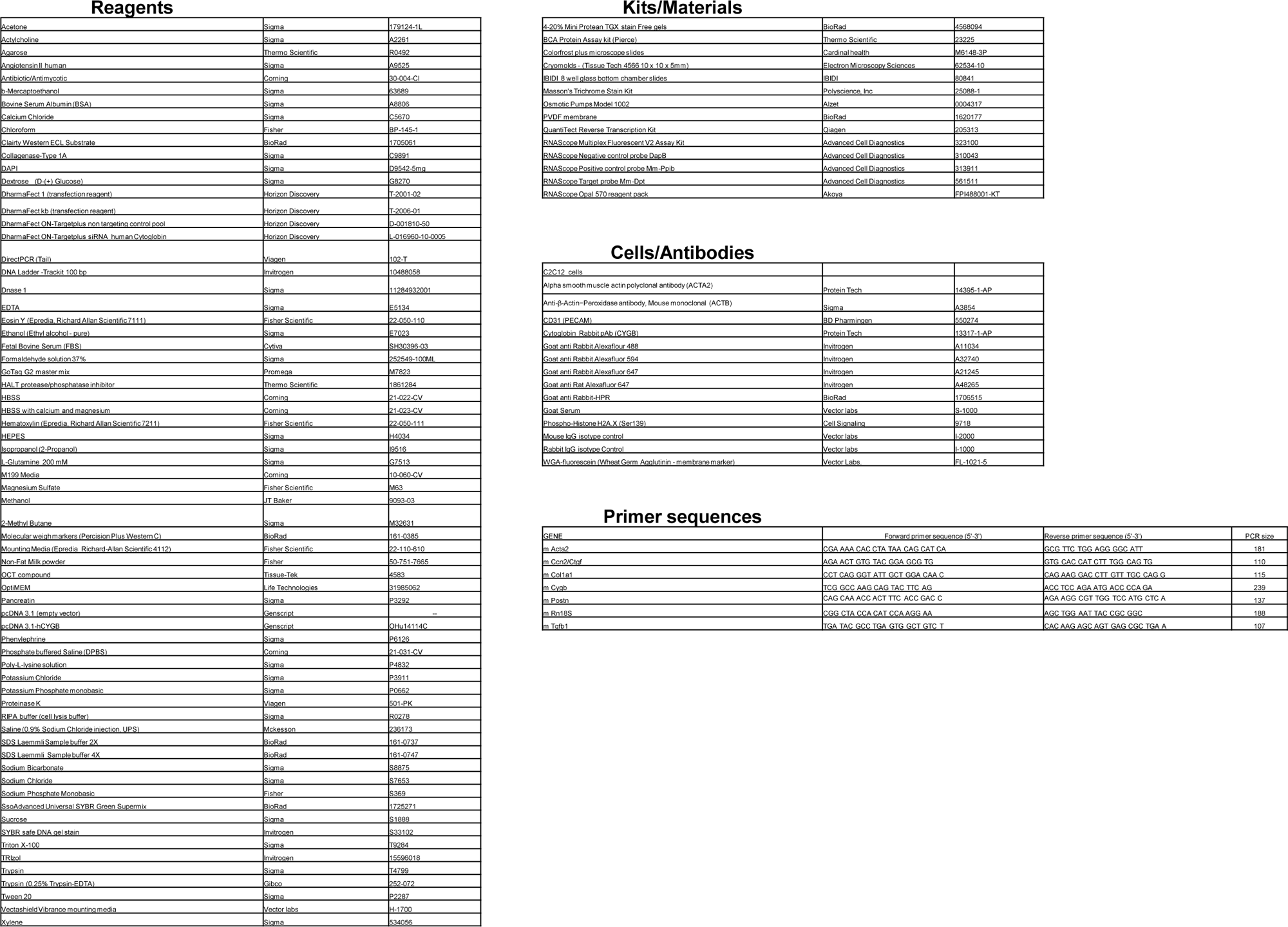
Supplies and reagents.

**Supplementary Figure S1.**
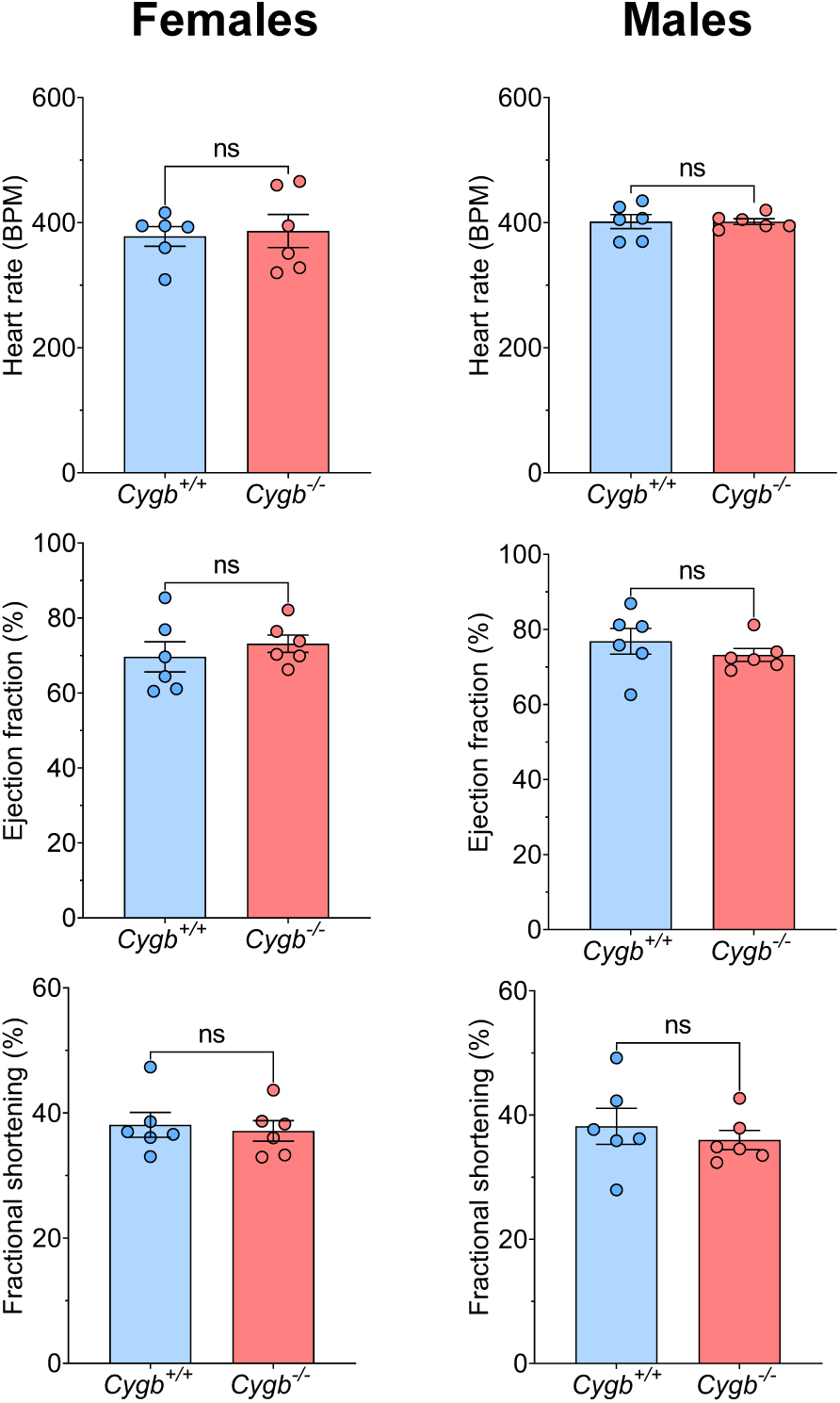
Global deletion of cytoglobin does not alter baseline cardiac function. Echocardiographic assessment of heart rate, ejection fraction, and fractional shortening was performed in female (left panels) and male (right panels) *Cygb^+/+^* and *Cygb^−/−^* mice under baseline conditions. No significant differences were observed between genotypes for any parameter in either sex (n = 6 per group). Data are presented as mean ± SEM, and statistical significance was determined by unpaired t-test; ns = not significant.

**Supplementary Figure S2.**
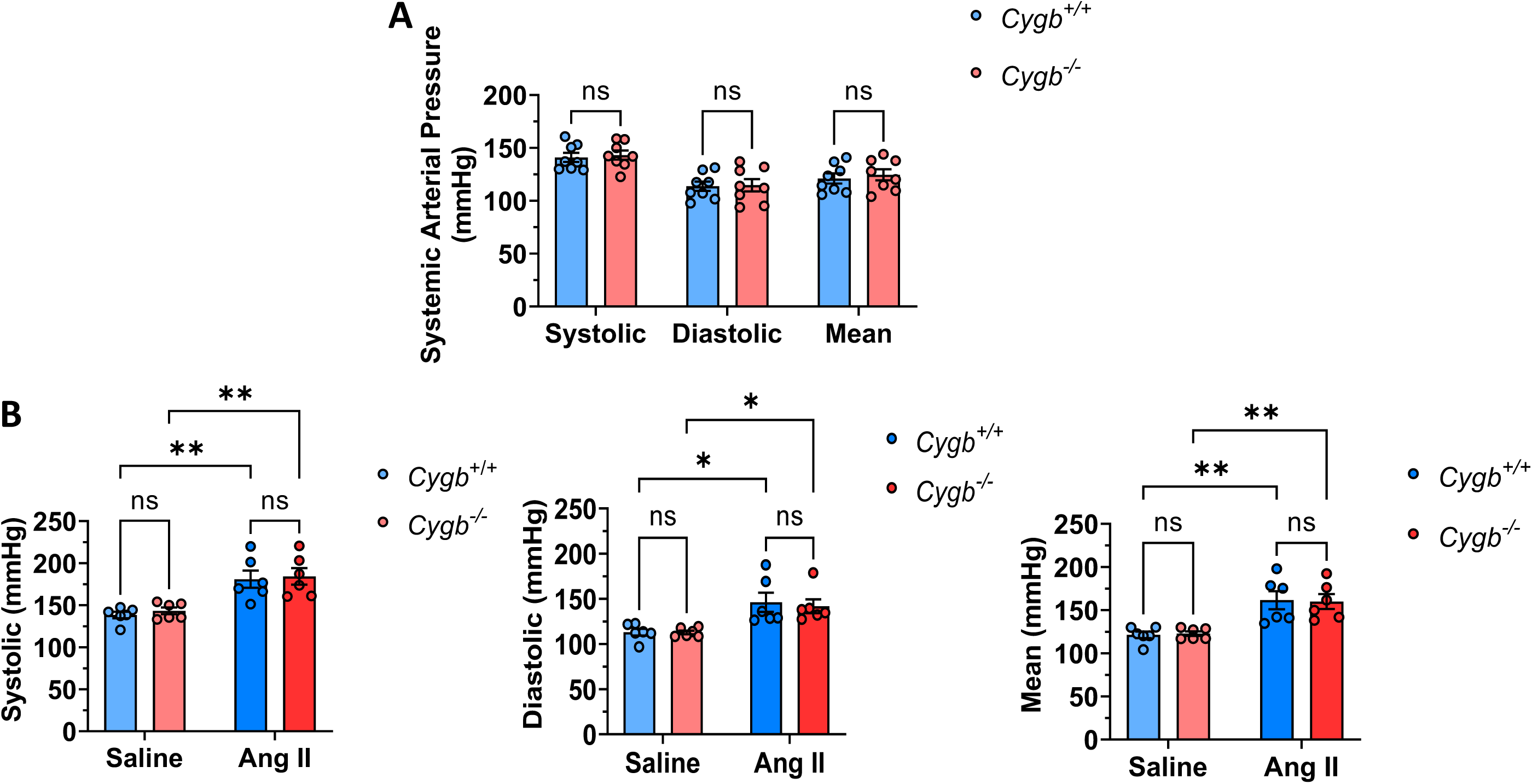
No change in blood pressure in female mice at baseline or following Angiotensin II infusion. **A:** baseline blood pressure in female *Cygb*^+/+^ and *Cygb*^−/−^ mice measured with a tail-cuff system. **B:** Female *Cygb*^+/+^ and *Cygb*^−/−^ mice (11–14 weeks old) were infused with saline or angiotensin II (Ang II; 1.5 mg/kg/day) for 2 weeks via subcutaneously implanted osmotic pumps. Systolic, diastolic, and mean arterial blood pressures were measured with a tail-cuff system. Data are presented as mean ± SEM, with statistical significance determined using two-way ANOVA. *P < 0.05, **P < 0.01, ns = not significant.

**Supplementary Figure S3.**
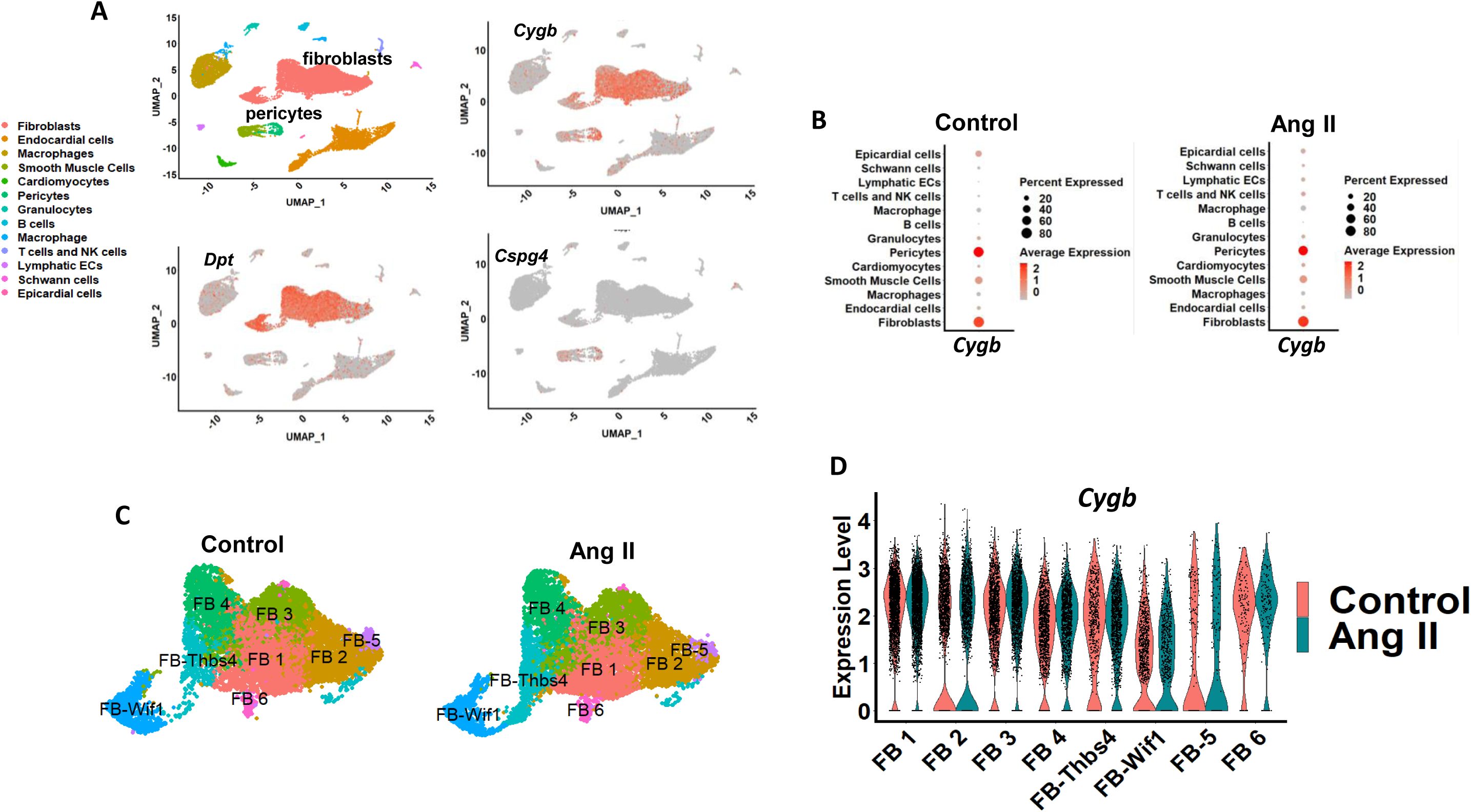
Cytoglobin expression in mouse hearts. **A:** Single-cell RNA-sequencing (scRNA-seq) data from publicly available mouse heart dataset (E-MTAB-8810) were reanalyzed to assess the expression pattern of *Cygb*, *Dpt*, *Cspg4* across major cardiac cell types. **B:** Dot plot indicates *Cygb* expression in control and Ang II treated hearts, enriched in fibroblasts and pericytes. Dot size represents the percentage of cells expressing Cygb, and color intensity indicates average expression level. **C:** UMAP plot of fibroblasts subpopulation in control and Ang II treated datasets. **D:** Violin plot showing *Cygb* expression levels in control and Ang II treated hearts.

**Supplementary Figure S4.**
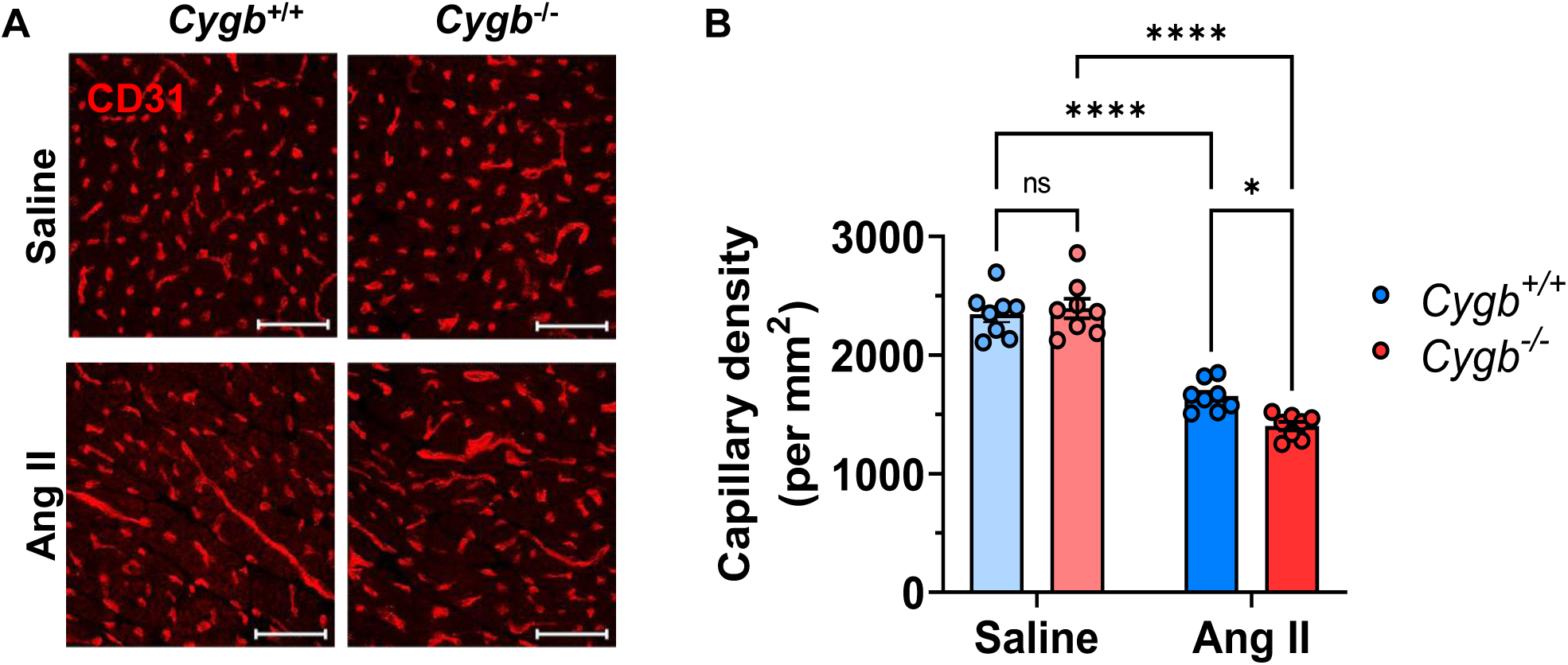
Cytoglobin deletion causes a small but significant decrease in capillary density. **A:** Representative left-ventricular sections from male *Cygb*^+/+^ and *Cygb*^−/−^ mice stained with CD31 to label capillaries. **B:** Quantitation of A. Scale bar = 100 µm. Each data point represents one mouse. Data are presented as mean ± SEM, with statistical significance determined using two-way ANOVA. *P < 0.05, ****P < 0.0001, and ns = not significant.

